# NanoVI: a Bayesian variational inference Nextflow pipeline for species-level taxonomic classification from full-length 16S rRNA Nanopore reads

**DOI:** 10.64898/2026.03.07.710315

**Authors:** Carolina Curiqueo, Francisco Fuentes-Santander, Juan A. Ugalde

## Abstract

**Summary:** NanoVI is a Nextflow pipeline for species-level taxonomic classification of full-length 16S rRNA Oxford Nanopore reads. Unlike existing tools that rely on expectation-maximization (EM) algorithms, NanoVI employs Bayesian variational inference with a Dirichlet–Categorical conjugate model, yielding abundance estimates accompanied by Bayesian 95% credible intervals that quantify estimation uncertainty, along with automatic shrinkage that suppresses spurious taxa. NanoVI integrates the Genome Taxonomy Database (GTDB) r226, providing phylogenetically consistent taxonomy while maintaining compatibility with NCBI-style databases. Benchmarked against a standardized mock community, NanoVI achieves species-detection metrics comparable to Emu, with 25–62% lower execution time and fewer false-positive assignments. Validation on 20 clinical vaginal microbiome samples confirms reproducibility against previously published Emu-based analyses.

**Availability and implementation:** NanoVI is implemented in Nextflow DSL2 with Docker containerization and is freely available at https://github.com/microbialds/NanoVI under an open-source license.

## 1 Introduction

The characterization of microbial communities through 16S ribosomal RNA gene sequencing has become the gold standard in microbial ecology and clinical diagnostics (Wood et al. 2019). Traditional approaches using short-read sequencing platforms like Illumina MiSeq, while offering high per-base accuracy (>99.9%), are fundamentally limited by their read length constraints of a few hundred base pairs (Caporaso et al. 2011). This limitation restricts taxonomic resolution to the genus level in most cases, as only specific variable regions of the 16S rRNA gene can be sequenced, missing crucial phylogenetic information contained in other regions (Johnson et al. 2019).

Oxford Nanopore Technologies (ONT) has enabled full-length 16S rRNA gene sequencing (∼1,500 bp), capturing all nine variable regions and providing species-level taxonomic resolution unattainable with short-read platforms (Woese and Fox 1977; Jain et al. 2016; Johnson et al. 2019). Recent advances in basecalling algorithms have substantially reduced Nanopore error rates, with high-accuracy models achieving median read qualities of Q18–Q22+ for 16S data (Wick et al. 2019; Aja-Macaya et al. 2025), supporting the growing suitability of ONT for microbiome analysis.

Several tools address the challenges of processing Nanopore 16S data. NanoCLUST (Rodríguez-Pérez et al. 2021) employs unsupervised clustering with UMAP and HDBSCAN, followed by BLAST-based classification. Emu (Curry et al. 2022) uses an EM algorithm to refine species-level abundance estimates from full-length reads iteratively. EPI2ME wf-16S v1.5.0 (Oxford Nanopore Technologies, 2025) supports multiple classifiers, including Minimap2 (Li, 2018) and Kraken2 (Wood et al. 2019). However, these tools face limitations: EM-based approaches treat species abundances as point estimates without quantifying uncertainty or implementing principled regularization against false positives; computational efficiency remains a bottleneck; and most tools rely on NCBI-style databases that may not reflect current phylogenetic understanding (Parks et al. 2018).

Here, we present NanoVI, a Nextflow pipeline that addresses these limitations through: (1) Bayesian variational inference based on coordinate ascent in a Dirichlet–Categorical model, providing Bayesian credible intervals for uncertainty quantification and automatic shrinkage; (2) integration of GTDB r226 (Parks et al. 2022) as the primary reference database; (3) systematic k-mer optimization for alignment; and (4) implementation in Nextflow DSL2 (Di Tommaso et al. 2017) with Docker containerization. We evaluate NanoVI on a standardized mock community and validate on 20 clinical vaginal microbiome samples from a recently published Chilean cohort (Oliva-Arancibia et al. 2025).

## 2 Methods

NanoVI is a modular pipeline implemented in Nextflow DSL2, containerized with Docker for reproducible execution across computing environments. The architecture comprises four functional modules: abundance estimation, database construction, taxonomy collapse, and output combination. The pipeline workflow is documented in https://github.com/microbialds/NanoVI.

### 2.1 Input Processing and Reference Database

The pipeline accepts raw ONT 16S reads in FASTQ format. Adapter trimming, quality filtering (minimum Q15), and length filtering (500–2,000 bp) are performed in a single pass using FastpLong v0.2.2 (Chen, 2023). NanoVI supports multiple reference databases, with primary support for the GTDB r226 database that contains 232,447 16S rRNA sequences representing 59,037 unique species (https://gtdb.ecogenomic.org/), and compatibility with NCBI-style databases. A build-database module constructs custom databases from user-provided FASTA files and automatically generates Minimap2 index files.

### 2.2 Alignment and Likelihood Estimation

Sequence alignment is performed using Minimap2 v2.24 (Li, 2018) with the map-ont preset and configurable k-mer sizes (default k=21, selected based on systematic optimization; Section 3); additionally, the maximum number of secondary alignments per read is set to N = 3 (versus N = 50 in Emu), limiting redundant alignments and contributing to reduced execution time. CIGAR strings are parsed to compute per-read alignment log-probabilities for each candidate reference: log_score(r, s) = [∑ log(pop) × countop] × (Lmax / Laln), where pop is the per-base probability for each CIGAR operation (match, mismatch, insertion, deletion) and the length normalization corrects for partial alignments.

### 2.3 Variational Inference Algorithm

NanoVI formulates abundance estimation as Bayesian inference in a Dirichlet–Categorical conjugate model, solved via mean-field Coordinate Ascent Variational Inference (CAVI) (Jordan et al. 1999; Blei et al. 2017). Species abundances π are assigned a symmetric Dirichlet prior with concentration α_0_ = 1 (distributing α_0_/S pseudo-counts uniformly across the S reference species), each read is assigned to a species via a Categorical distribution, and observed alignment data provide species-specific likelihoods. The intractable posterior is approximated via the factored family q(π, z) = q(π) ∏r q(zr), with CAVI iteratively updating read-to-species responsibilities and Dirichlet parameters. The use of digamma functions in the updates introduces Bayesian shrinkage: species with weak alignment evidence are automatically downweighted, reducing false-positive detections (Parks et al. 2022). Upon convergence, posterior means provide abundance point estimates, and 95% credible intervals—derived analytically from the Beta marginals π□ ∼ Beta(α□, Σ□ α□ − α□)—quantify the uncertainty of each species estimate. An outer pruning loop removes species below a data-adaptive threshold and re-runs CAVI until convergence.

### 2.4 Output and Post-processing

NanoVI produces per-sample relative abundance tables with Bayesian 95% credible intervals reflecting estimation uncertainty for each taxon, along with estimated read counts, across seven hierarchical taxonomic ranks (species, genus, family, order, class, phylum, and superkingdom). The write_output module generates tab-separated results including relative abundances, estimated read counts, and full taxonomic lineage annotations per taxon. The collapse-taxonomy module aggregates per-sample abundance estimates at any configurable taxonomic rank. The combine-outputs module merges sample-specific tables into unified abundance. It counts matrices with taxa as rows and samples as columns, ready for import into popular microbiome analysis frameworks such as phyloseq (R). All pipeline parameters are user-configurable through command-line flags or configuration files. Key parameters include filtering thresholds (--min-length, --max-length, --min-quality), alignment configuration (--kmer-size, --type), computational resources (--threads), and taxonomic resolution (--rank). Based on our systematic evaluation (Section 3.1, Figure S1), k=21 was selected as the recommended default parameter, providing an optimal balance between runtime (6.55 min), memory usage (14.9 GB), and taxonomic accuracy.

All Python scripts were executed in a containerized environment running Python v3.8, using the following libraries: NumPy v1.21.6, Pandas v1.3.5, PySAM v0.19.1, and Biopython v1.79.

## 3 Results and Discussion

NanoVI was evaluated on the Zymo Research mock community dataset (SRR23636353). The theoretical 16S rRNA gene abundance was corrected from the equal genomic DNA composition (12% each) by genome size and 16S rRNA gene copy number: *L. fermentum* (0.184), *B. subtilis* (0.174), *S. aureus* (0.155), *L. monocytogenes* (0.141), *S. enterica* (0.104), *E. coli* (0.101), *E. faecalis* (0.099), and *P. aeruginosa* (0.042). These values served as the expected composition for all accuracy metrics. Clinical validation was performed on 20 clinical vaginal microbiome samples (Oliva-Arancibia et al. 2025). For reproducibility, three independent subsampled replicates (80% of reads) were generated per condition using seqtk v1.4 (Li, 2023). All analyses were performed on a workstation with 14 CPU threads and 128 GB RAM. Evaluation metrics included Bray-Curtis dissimilarity, Jensen-Shannon divergence, L1 distance, precision, recall, F1, and AUPRC.

### 3.1 k-mer Optimization and Computational Performance

To evaluate the impact of k-mer size on computational performance, we systematically tested k-mer sizes ranging from 15 to 28 using the mock community dataset (SRR23636353), running three independent subsampled replicates (80% of reads) per k-mer size (Figure S1, Supplementary Table 1). Execution time decreased substantially with increasing k-mer size, from 15.60 min at k=15 to 3.87 min at k=28, representing a 4-fold reduction. Peak RAM usage decreased more gradually, from 16.1 GB at k=15 to 14.2 GB at k=28, remaining within the capacity of standard workstations throughout. The standard deviations across replicates were uniformly negligible (≤0.04 min for execution time and ≤0.04 GB for peak RAM), confirming pipeline reproducibility.

Taxonomic recovery was robust across the entire tested range (Figure 1). NanoVI successfully detected all eight expected species from the Zymo mock community at every k value, demonstrating that larger k-mers do not compromise taxonomic completeness. Notably, the GTDB r226 database resolved multiple species within the Bacillus subtilis complex not distinguishable in NCBI-based databases. Low-abundance taxa originally present in mock communities (e.g., *B. subtilis, P. aeruginosa*) were consistently detected, confirming the sensitivity of the CAVI approach. Based on the trade-off between execution time, memory usage, and taxonomic completeness, we selected k=21 as the recommended default for NanoVI.

**Figure 1.**
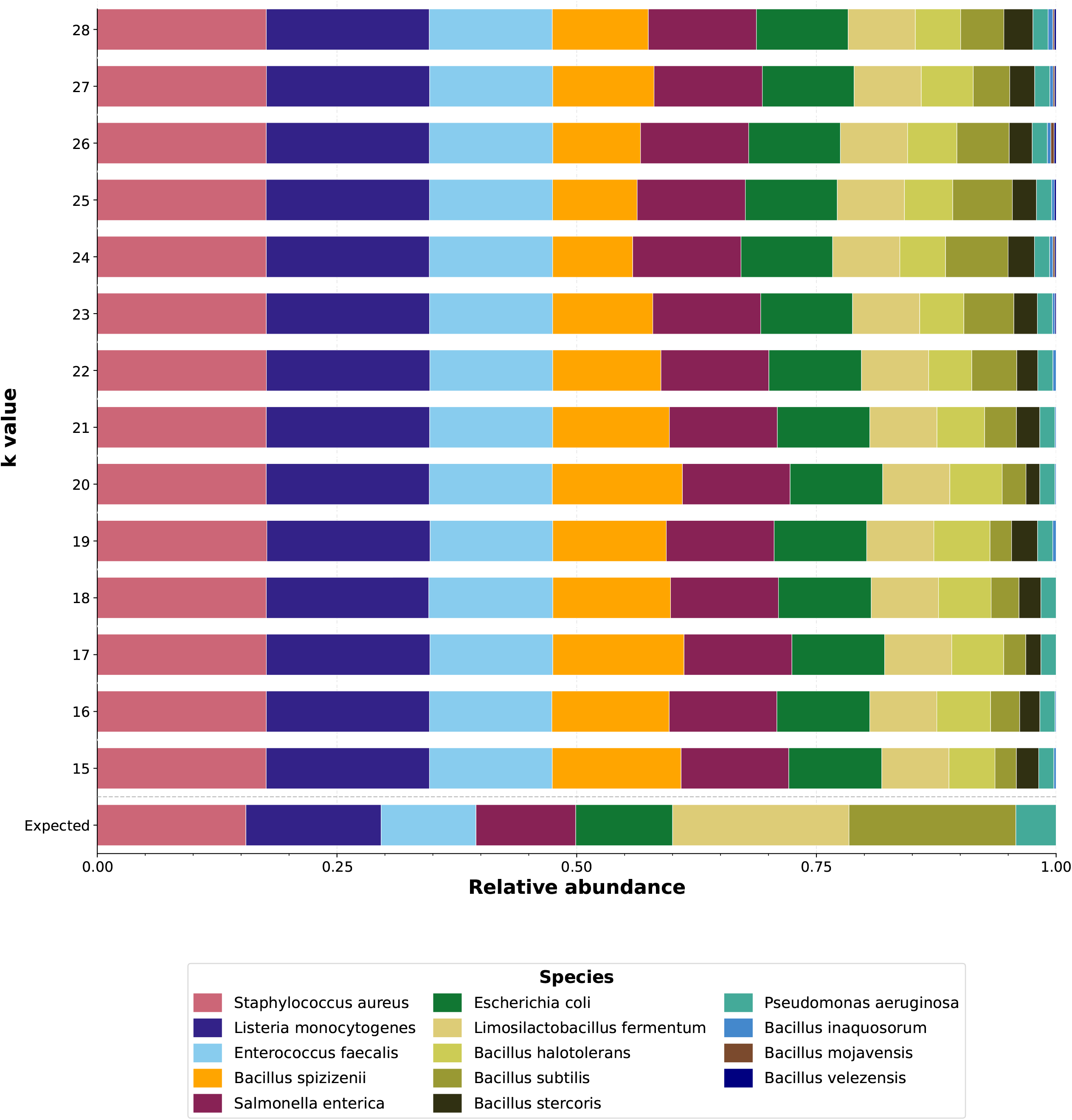
Taxonomic composition of mock communities classified by NanoVI across k-mer sizes using GTDB r226. Each bar represents a separate NanoVI run at the indicated k-mer value (k=15 to k=28, y-axis). The x-axis shows relative abundance (0–1). Colors indicate species as listed in the legend.

A central methodological contribution of NanoVI is the replacement of EM-based point estimation with Bayesian variational inference, which provides posterior credible intervals that enable researchers to flag low-confidence detections and account for uncertainty in comparative analyses. This probabilistic framework, combined with the computational gains described above, is particularly relevant for clinical diagnostics requiring both rapid turnaround and robust uncertainty quantification.

### 3.2 Comparative Analysis

To isolate the effect of the inference algorithm from database choice, we first compared NanoVI and Emu using the same Emu NCBI-style reference database on the Zymo mock community (Figure 2). NanoVI at both k=15 and k=21 produced taxonomic profiles highly concordant with those generated by Emu (Figure 2a), with all eight major taxa detected at consistent relative abundances. Community-level distance metrics and species detection performance confirmed this concordance (Figures 2b–2c): both NanoVI configurations achieved dissimilarity and detection scores comparable to Emu, with precision, recall, F1, and AUPRC all approaching 1.0. Minor differences included the detection of *Laceyella sacchari* and *Shigella dysenteriae* at very low abundances by Emu and NanoVI at k=21, respectively, likely reflecting variation in low-confidence read assignments and the Bayesian shrinkage effect of the Dirichlet prior. Critically, NanoVI achieved these comparable accuracy levels with substantially reduced computational time: a 25% reduction at k=15 (12.21 ± 0.03 min vs. 16.44 ± 0.09 min) and a 62% reduction at k=21 (6.55 ± 0.02 min vs. 16.44 ± 0.09 min), under identical hardware conditions (Table 1).

**Figure 2.**
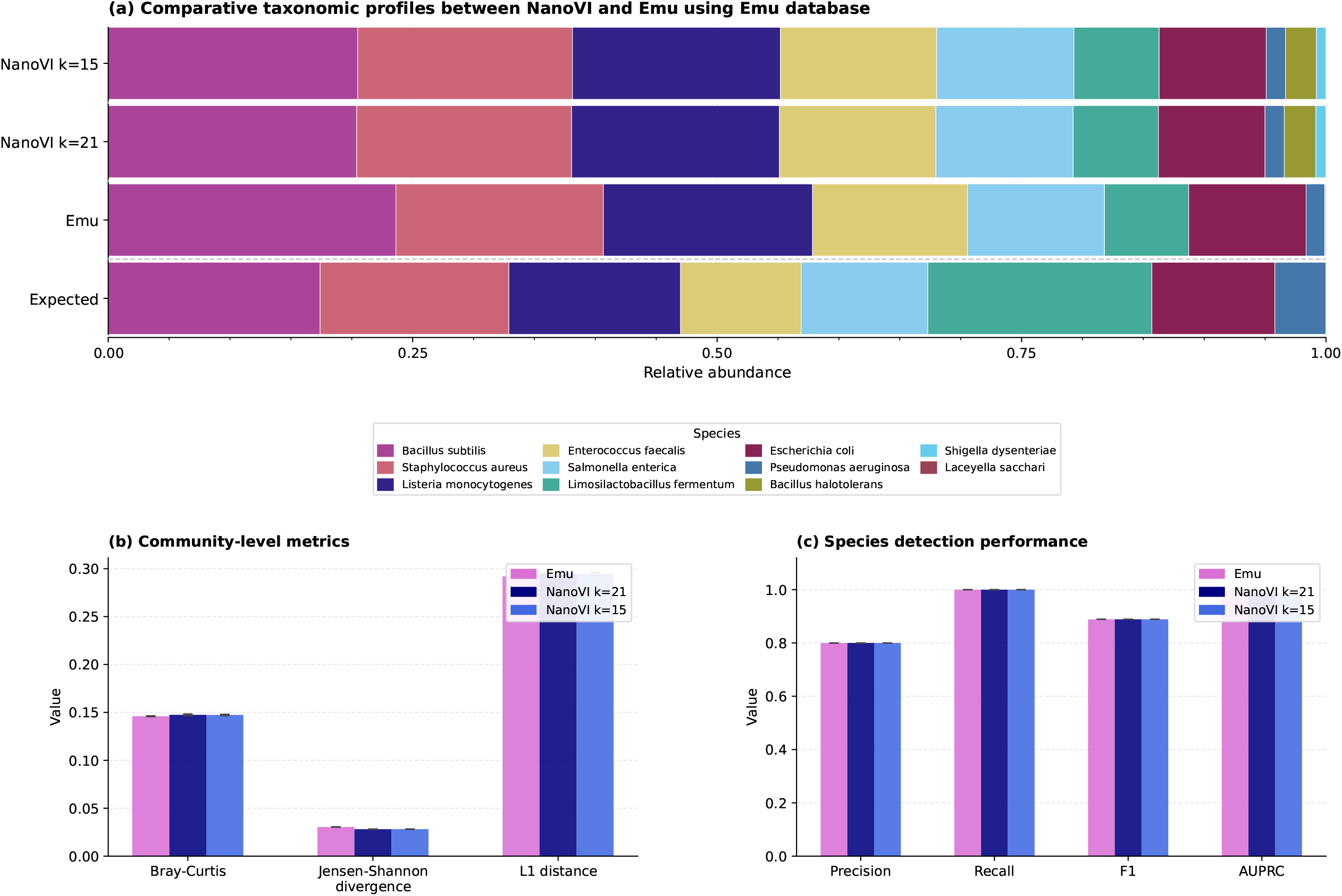
Taxonomic profiles and accuracy metrics for NanoVI and Emu on a mock community using the Emu NCBI-style reference database. (a) Stacked bar charts showing relative abundance (x-axis, 0–1) for each tool (Emu, NanoVI k=21, NanoVI k=15; y-axis). Colors indicate species as listed in the legend. (b) Community-level dissimilarity metrics (Bray-Curtis dissimilarity, Jensen-Shannon divergence, and L1 distance; y-axis) computed against the expected composition, for each tool (x-axis). (c) Species detection performance metrics (Precision, Recall, F1, and AUPRC; x-axis, range 0–1) for each tool (colored bars). All NanoVI values represent means across three independent replicates (80% read subsampling; SD not visible at this scale).

**Table 1.**
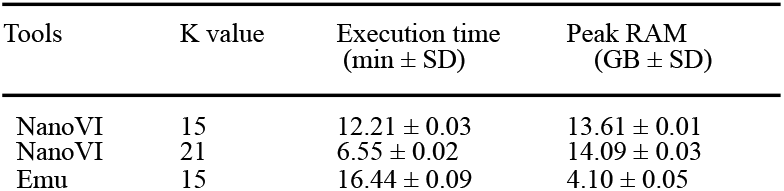
Execution time and peak RAM usage for NanoVI and Emu on a mock community. All runs used the Emu NCBI-style reference database on identical hardware (14 CPU threads, 128 GB RAM). NanoVI and Emu values are means ± SD across three independent replicates (80% random read subsampling). The k value corresponds to the k-mer length used in Minimap2.

We then extended the comparison to a broader set of tools: Emu (default NCBI-style database), NanoCLUST, and EPI2ME wf-16S, each using their default databases on the same mock community dataset (Figure 3). NanoVI and Emu produced highly similar taxonomic profiles, successfully recovering all major taxa with appropriate relative abundances (Figure 3a). In contrast, NanoCLUST and EPI2ME assigned a substantially larger fraction of reads to an “Other” category and failed to detect several key species, including *Staphylococcus aureus* and *Listeria monocytogenes*. These differences were reflected in the quantitative metrics (Figures 3b–3c): NanoVI achieved community-level dissimilarity and species detection scores comparable to Emu and substantially superior to NanoCLUST and EPI2ME. Although NanoCLUST (3.77 ± 0.38 min) and EPI2ME (3.46 ± 0.12 min) were faster than NanoVI (6.55 ± 0.02 min; Table 2), this speed advantage came at a substantial cost in taxonomic accuracy, underscoring that NanoVI offers a favorable trade-off between computational efficiency and classification performance.

**Table 2.** Execution time and peak RAM usage for NanoVI, Emu, NanoCLUST, and EPI2ME wf-16S on a mock community (SRR23636353). Each run with its default reference database on identical hardware (14 CPU threads, 128 GB RAM). NanoVI, Emu NanoCLUST, and EPI2ME values are mean ± SD across three independent replicates (80% random read subsampling).

| Tools | Database | K value | Execution time (min ± SD) | Peak RAM (GB ± SD) |
| --- | --- | --- | --- | --- |
| EPI2ME | Default | - | 3.46 ± 0.12 | 11.32 ± 0.02 |
| NanoCLUST | Default | - | 3.77 ± 0.38 | 9.03 ± 0.06 |
| NanoVI | GTDB r226 | 21 | 6.55 ± 0.02 | 14.09 ± 0.03 |
| Emu | Emu | 15 | 16.44 ± 0.09 | 4.10 ± 0.05 |

**Figure 3.**
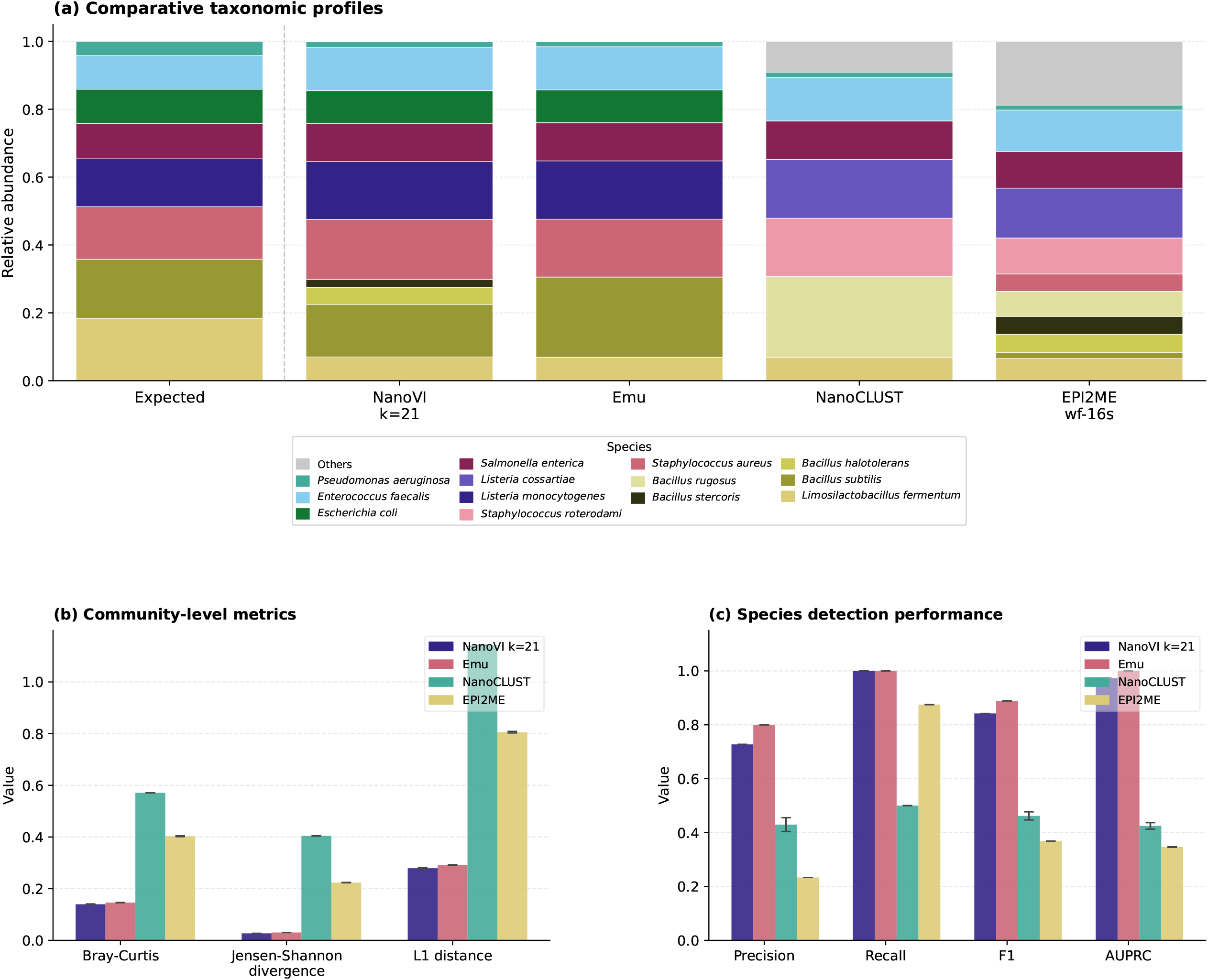
Taxonomic profiles and accuracy metrics for four 16S classification tools on a mock community. (a) Stacked bar charts showing relative abundance (x-axis, 0–1) for NanoVI k=21 (GTDB r226), Emu, NanoCLUST, and EPI2ME wf-16S (y-axis). Taxa below 1% relative abundance are grouped into “Other.” Colors indicate species as listed in the legend. (b) Community-level dissimilarity metrics (Bray-Curtis dissimilarity, Jensen-Shannon divergence, and L1 distance; y-axis, range 0–1) computed against the expected composition, for each tool (x-axis). (c) Species detection performance metrics (Precision, Recall, F1, and AUPRC; x-axis, range 0–1) for each tool (colored bars).

The speed gains are attributable to two complementary factors: k-mer optimization (Section 3.1) and a reduced secondary alignment limit (N = 3 versus N = 50 in Emu), which together limit redundant computations without compromising taxonomic accuracy.

### 3.3 Clinical Validation

To evaluate NanoVI’s performance on real clinical samples, we applied the pipeline to full-length 16S rRNA Nanopore sequencing data from 20 vaginal microbiome samples (M1–M20) previously published by Oliva-Arancibia et al. 2025, a study characterizing the vaginal microbiota of Chilean women using self-sampling and ONT sequencing. The original study employed Emu with the Emu NCBI-style reference database. Here, we re-analyzed the same dataset with NanoVI using two reference databases: GTDB r226 and the Emu reference database, to assess reproducibility and evaluate the added resolution offered by GTDB (Figure 4). This cross-validation against a published reference analysis constitutes a practical test involving real samples with complex community compositions. Moreover, compatibility with existing Emu-based datasets enables retrospective re-analysis of published cohorts with Bayesian uncertainty quantification and updated taxonomic nomenclature.

**Figure 4.**
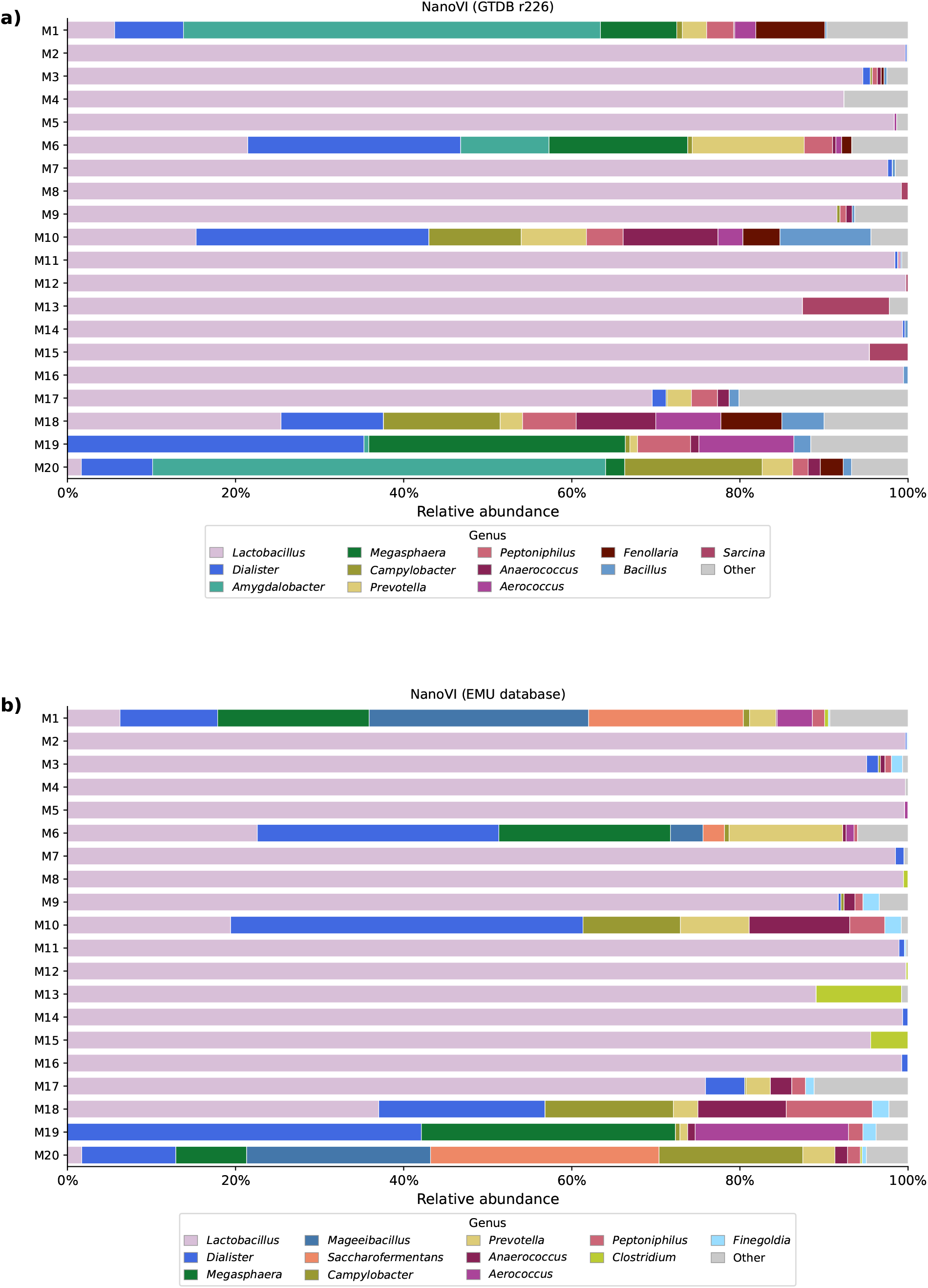
Genus-level taxonomic profiles of vaginal microbiome samples classified by NanoVI. (a) NanoVI with GTDB r226. (b) NanoVI with the Emu NCBI-style reference database. In both panels, the x-axis shows relative abundance (0–100%), and colors indicate genera as listed in the legend. Only the 12 most abundant genera (by mean relative abundance across all samples) are shown individually; remaining genera are grouped into “Other.”

Both NanoVI configurations consistently reproduced the community-level structure originally reported. *Lactobacillus* dominated the majority of samples, in full agreement with the original Emu-based results, while samples M1, M6, M10, and M18 exhibited polymicrobial profiles with co-dominant *Dialister, Megasphaera, Prevotella, Anaerococcus, Peptoniphilus, Aerococcus*, and *Campylobacter*, consistent with the dysbiotic profiles previously described. NanoVI with the Emu database (Figure 4b) produced profiles closely mirroring the original publication, whereas NanoVI with GTDB r226 (Figure 4a) produced concordant community profiles with some taxonomic reassignments reflecting the phylogenetically consistent classification system of GTDB.

The integration of GTDB r226 as the primary reference database provides a phylogenetically consistent taxonomy that resolves polyphyletic genera persisting in NCBI. A notable example is *Clostridium*, which in NCBI groups phylogenetically distant lineages under a single genus name. In our vaginal microbiome analysis, reads classified as *Clostridium* under NCBI were reassigned to *Sarcina* (https://gtdb.ecogenomic.org/), reflecting the phylogenetically consistent reclassification of polyphyletic genera in GTDB (Parks et al. 2022; Cruz-Morales et al. 2019). This illustrates how GTDB resolves taxonomic inconsistencies that NCBI-based databases retain, although cross-database comparisons require explicit taxonomic mapping.

### 3.4 Limitations and Future Directions

Despite its advantages, NanoVI has limitations. Evaluation has also been limited primarily to bacterial communities, and validation on archaeal or eukaryotic microorganisms would be valuable for users working with diverse microbial communities.

Future development could explore GPU acceleration, streaming CAVI implementations, or sparse matrix approximations for extremely large-scale analyses. Integration of phylogenetic diversity metrics (e.g., weighted UniFrac) and functional prediction capabilities would further expand NanoVI’s analytical scope. Automated database update mechanisms would ensure users maintain access to the most current GTDB releases without manual intervention.

## 4 Conclusions

NanoVI combines Bayesian variational inference with GTDB r226 integration to classify species-level 16S rRNA from Nanopore data, achieving 25–62% reductions in execution time relative to Emu, with comparable accuracy and fewer false positives than other tools. Current limitations include benchmarking primarily on low-to-moderate diversity communities and the dense matrix representation that may become memory-prohibitive for very large reference databases. Future development will address GPU acceleration, sparse-matrix approximations, and the integration of phylogenetic diversity metrics.

## Supplementary data

Supplementary data are available at *Bioinformatics Advances* online.

## Data availability

The Zymo mock community data are available in the NCBI SRA (SRR23636353). NanoVI source code and documentation are available at https://github.com/microbialds/NanoVI.

## Supplementary data

**Supplementary Figure S1. NanoVI computational performance across k-mer sizes 15–28.** (a) Total execution time (min, left y-axis; blue circles, solid line) and peak RAM usage (GB, right y-axis; red squares, dashed line) as a function of k-mer size (x-axis, 15–28). (b) Execution time (min, y-axis) per pipeline step (colors as listed in the legend) for each k-mer value (x-axis).

