## Supplemental Figure 1 for "NanoVI: a Bayesian variational inference Nextflow pipeline for species-level taxonomic classification from full-length 16S rRNA Nanopore reads"

**Supplementary Figure 1. NanoVI computational performance across k-mer sizes**

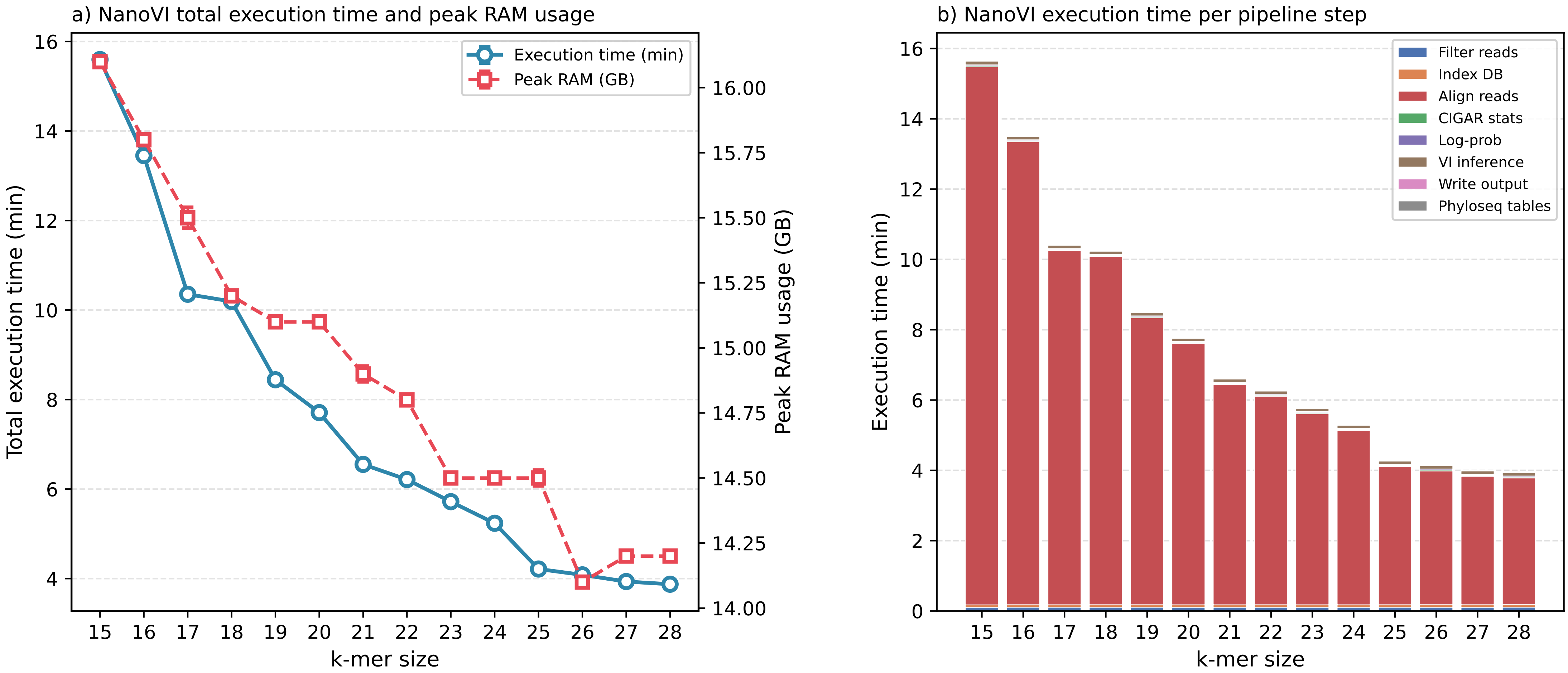


**Supplementary Table 1. NanoVI computational performance for each k-mer value**

| k value | Execution time (min) ± SD | Peak RAM (GB) ± SD |
|---|---|---|
| **15** | 15.60 ± 0.01 | 16.1 ± 0.01 |
| **16** | 13.45 ± 0.02 | 15.8 ± 0.01 |
| **17** | 10.35 ± 0.02 | 15.5 ± 0.04 |
| **18** | 10.19 ± 0.04 | 15.2 ± 0.02 |
| **19** | 8.44 ± 0.00 | 15.1 ± 0.01 |
| **20** | 7.71 ± 0.02 | 15.1 ± 0.01 |
| **21** | 6.55 ± 0.02 | 14.9 ± 0.03 |
| **22** | 6.21 ± 0.02 | 14.8 ± 0.01 |
| **23** | 5.72 ± 0.01 | 14.5 ± 0.01 |
| **24** | 5.24 ± 0.04 | 14.5 ± 0.02 |
| **25** | 4.21 ± 0.00 | 14.5 ± 0.03 |
| **26** | 4.08 ± 0.01 | 14.1 ± 0.01 |
| **27** | 3.93 ± 0.01 | 14.2 ± 0.01 |
| **28** | 3.87 ± 0.01 | 14.2 ± 0.02 |
